# Effectiveness of live attenuated monovalent human rotavirus vaccination in rural Ecuador, 2008-2013

**DOI:** 10.1101/482893

**Authors:** Alicia N. M. Kraay, Edward L. Ionides, Gwenyth O. Lee, William F. Cevallos Trujillo, Joseph N.S. Eisenberg

**Affiliations:** Department of Epidemiology, University of Michigan-Ann Arbor, 1415 Washington Heights Ann Arbor, MI 48109-2029; Department of Statistics, University of Michigan-Ann Arbor, 1085 South University Ave Ann Arbor, MI 48109-1107; Centro de Biomedicina. Facultad de Ciencias Médicas, Universidad Central del Ecuador, Quito, Ecuador

**Keywords:** Rotarix, rotavirus, vaccine effectiveness, Ecuador, rural

## Abstract

**Background:** While live attenuated monovalent human rotavirus vaccine (Rotarix) efficacy has been characterized through randomized studies, its effectiveness, especially in non-clinical settings, is unclear. In this study, we estimate direct, indirect, and overall effectiveness of Rotarix vaccination.

**Methods:** We analyze 29 months of all-cause diarrhea surveillance from a child cohort (n=376) and ten years of serial population-based case-control lab-confirmed rotavirus data (n=2489) from rural Ecuador during which Rotarix vaccination was introduced. We estimate: 1) the direct effect of vaccination from a cohort of children born from 2008-2013 using Cox regression to compare time to first all-cause diarrhea case by vaccine status; and 2) the overall effect on all-cause diarrheal and symptomatic and asymptomatic rotavirus infection for all age groups, including indirect effects on adults, from the case-control data using weighted logistic regression.

**Results:** Rotarix vaccination provided direct protection against all-cause diarrhea among children 0.5 - 2 years (All-cause diarrhea reduction for receipt of 2 doses of Rotarix=57.1%, 95% CI: 16.6, 77.9%). Overall effectiveness against rotavirus infection was strong (Exposure to 100% coverage of Rotarix vaccination was associated with an 85.5% reduction, 95% CI: 61.1-94.6%) compared to 0% coverage. Indirect effects were observed among older, vaccine-ineligible children and adults (84.5% reduction, 95% CI: 48.2-95.4%). Vaccine effectiveness was high against both symptomatic (48.3% reduction,95% CI: 0.03-73.1%) and asymptomatic infection (90.1% reduction, 95% CI: 56.9-97.7%).

**Conclusions:** Rotarix vaccination suppresses overall transmission. It is highly effective among children in a rural community setting and provides population-level benefits through indirect protection among adults.

---

Rotavirus is a major cause of severe diarrhea worldwide, particularly in young children [1–3]. In 2006 the Rotarix and Rotateq vaccines were approved with the objective of reducing severe rotavirus [3,4]. While both vaccines have shown similar effectiveness, Rotarix is used in Ecuador and in most lower-and-middle income countries (LMICs) because of its better cost effectiveness, lower dose requirements, and thermostability [5–7]. In general, Rotarix vaccination is thought to be most effective against severe rotavirus infection and decreases the burden of severe diarrhea in populations where it is introduced [8–14]. However, most studies on rotavirus vaccine performance have come from clinical populations; data on how the vaccine performs against milder disease and asymptomatic infections especially are lacking [5]. Moreover, many of the prior studies have focused on large cities [8–11]; data on how the vaccine performs in community settings, particularly in rural settings, are needed [5]. Here we examine Rotarix effectiveness on rotavirus infections (including asymptomatic infections) and all-cause diarrhea in a rural community setting in Ecuador.

In addition to demonstrating efficacy against severe rotavirus among young children, some studies have also shown reductions in rotavirus infections in unvaccinated populations [15–17], suggesting that rotavirus vaccination may reduce the transmission rate. However, most evidence of this effect comes from high income countries [15, 16]. There is a need for data to determine if vaccination also reduces transmission rates in LMICs, which could increase the overall total impact of vaccination in such settings [5, 17]. It may also be easier to interrupt transmission in rural regions of LMICs because some are partially protected from illness due to their relative isolation [18]. In this context, rotavirus transmission patterns are driven by periodic reintroductions by older children and adults, who are not age-eligible to be vaccinated [19]. Therefore, indirect protection might help maintain vaccine effects over time by reducing community susceptibility to periodic reintroduction.

Our objective is to determine if Rotarix implementation has reduced rotavirus infection, including asymptomatic infection, and all-cause diarrhea in a rural region of Ecuador with high levels of endemic diarrhea. We use these results to quantify 1) indirect effects of vaccination on un-vaccinated populations and 2) the impact of vaccination on asymptomatic rotavirus infection.

## Methods

### Data

This analysis uses data collected for a study on diarrheal disease in rural Ecuador over a ten-year period (from 2003 to 2013, Figure 1) [18]. This included: yearly census information, active all-cause diarrhea surveillance, and periodic case control data. We also collected vaccine records for children born in study villages between 2008 and 2013. Rotarix was introduced in Ecuador in 2007, but did not begin being administered in the study region until late August of 2008 [20]. Using date of birth, community of residence, and name, we linked these records to individuals in our larger study (see S1 for more details). All protocols were approved by Institutional Review Boards at University of Michigan and Universidad San Francisco de Quito.

**Figure 1.**
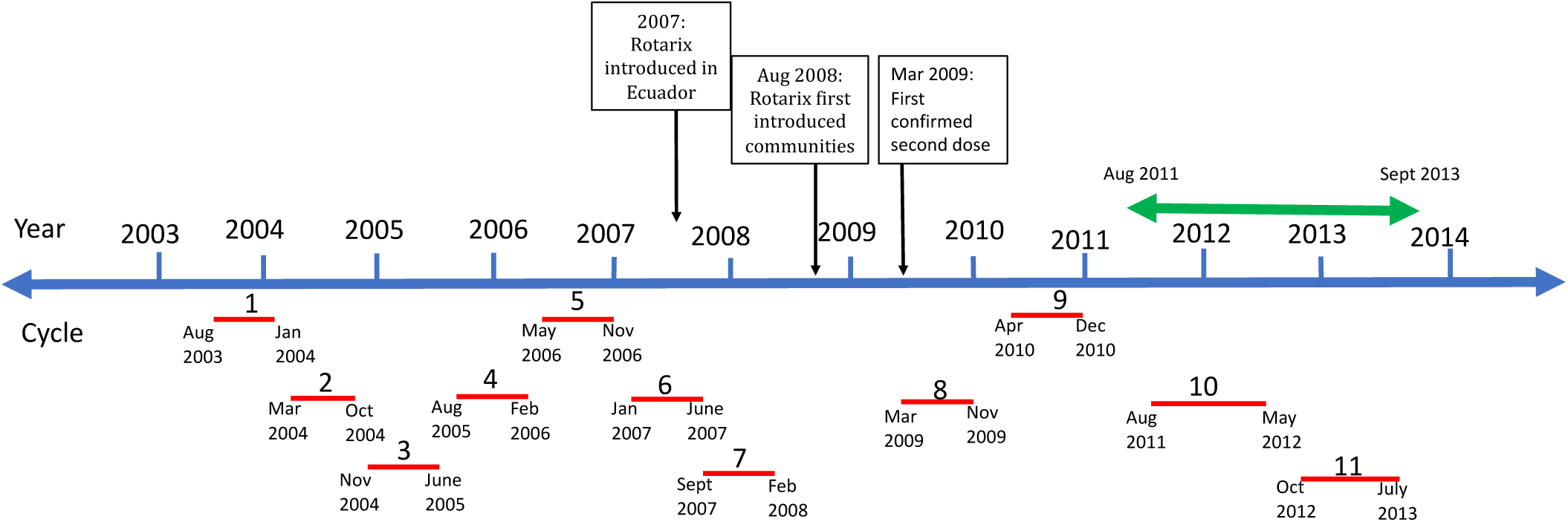
Data collection schedule, where each calendar year is shown in blue. Throughout the ten-year study period, we conducted 11 serial, population-based, case control studies (timing shown in red). The population-based study design means that each individual in the community had a known probability of sampling. We conducted an active surveillance study between August 2011 and September 2013 (timing shown in green), and a census in each cycle prior to the start of the case control study. Rotarix was introduced in Ecuador in 2007 but was not introduced in our study communities until late 2008.

#### Outcome data

Between August 2011 and December 2013 we collected diarrheal surveillance data from all households every two weeks. This all-cause diarrhea surveillance did not include any testing for rotavirus status. Approximately once every 9 months from 2003 to 2013, staff visited each community and conducted a two week, prospective, population-based case-control study for symptomatic diarrhea. Each case control study cycle occurred over an approximate 6-month study period and included communities sampled in the main rainy (December-May) and dry season (June-November) [21]. The order of sampling for each community rotated over time, such that data are available from both seasons for all communities (see Figure 1). In total, we have 11 cycles of case control data for fifteen communities. During each 14-day period, cases of diarrhea were identified prospectively. A case in the case control study was defined by having three or more loose stools in the last 24 hours, irrespective of the pathogen causing that diarrhea. For the first seven study cycles, we collected one household control and two community controls per case. For cycles 8-11, a random sample of 10% of the community population was sampled as controls at baseline. We obtained stool samples from each case and control and these samples were tested for rotavirus (using the EIA kit, RIDA Quick Rotavirus; R-Biopharm, Darmstadt, Germany). While we did not collect data on severity of diarrheal disease episodes for either the active surveillance or the case-control study, it is likely that cases are milder on average than in other studies that have focused on medically attended diarrhea.

#### Covariate data

We quantified the remoteness of each village based on the cost and travel time required to reach Borbón, the largest town in the region when the study began (2003) [18]. Both cost and travel time were normalized to fall between 0 and 1, and the combined remoteness score combined information from both variables. This measure was subsequently categorized into 3 groups: ‘close,’ ‘intermediate,’ and ‘far.’ Most ‘close’ communities were connected to Borbón by a road at the start of the study, requiring less than 30 minutes to reach the town by car, whereas the ‘far’ communities lacked road access and were several hours from the town by motorized canoe. Intermediate communities were moderately remote, and access to Borbón was intermediate between these two extremes. As road development was ongoing throughout the study period, the absolute time and cost to reach Borbón decreased over time, but the relative frequency of travel remained similar [19].

For the surveillance analysis, we categorized participant age into 6 months-2 years and ≥2 years based on age at entry into the cohort. For the case control analysis, we used 3 age groups: <1, 1-5, and ≥5 years.

We adjusted for household socioeconomic status using highest household education (highest number of years of schooling reported by any household member) and number of children in the household. For the surveillance, household characteristics were calculated from the census nearest to the child’s date of birth. For case control analyses, socioeconomic indicators were taken from the census nearest to the case-control cycle. For the surveillance analysis, we also adjusted for BCG vaccination (yes/no) because it is administered at the same time as rotavirus and therefore accounts for health seeking behavior and has been shown to have beneficial non-specific effects [22].

### Analytic approach

To assess the effect of Rotarix vaccination in our study site, we conducted a two-part analysis (see table 1), using the convention of Halloran and Hudgens to estimate vaccine effectiveness (VE; see S2) [23]. First, we assessed the association between community vaccine coverage (proportion of children with vaccine records who received their second dose of Rotarix during a given case-control cycle) on both rotavirus infection (including asymptomatic infection) and all-cause diarrhea, using ten years of case-control data. Second, we assessed the direct effect of vaccination on all-cause diarrhea illness for children under five years using 29 months of active surveillance data. More methodological details about the analysis are found in S3-S4.

**Table 1:**
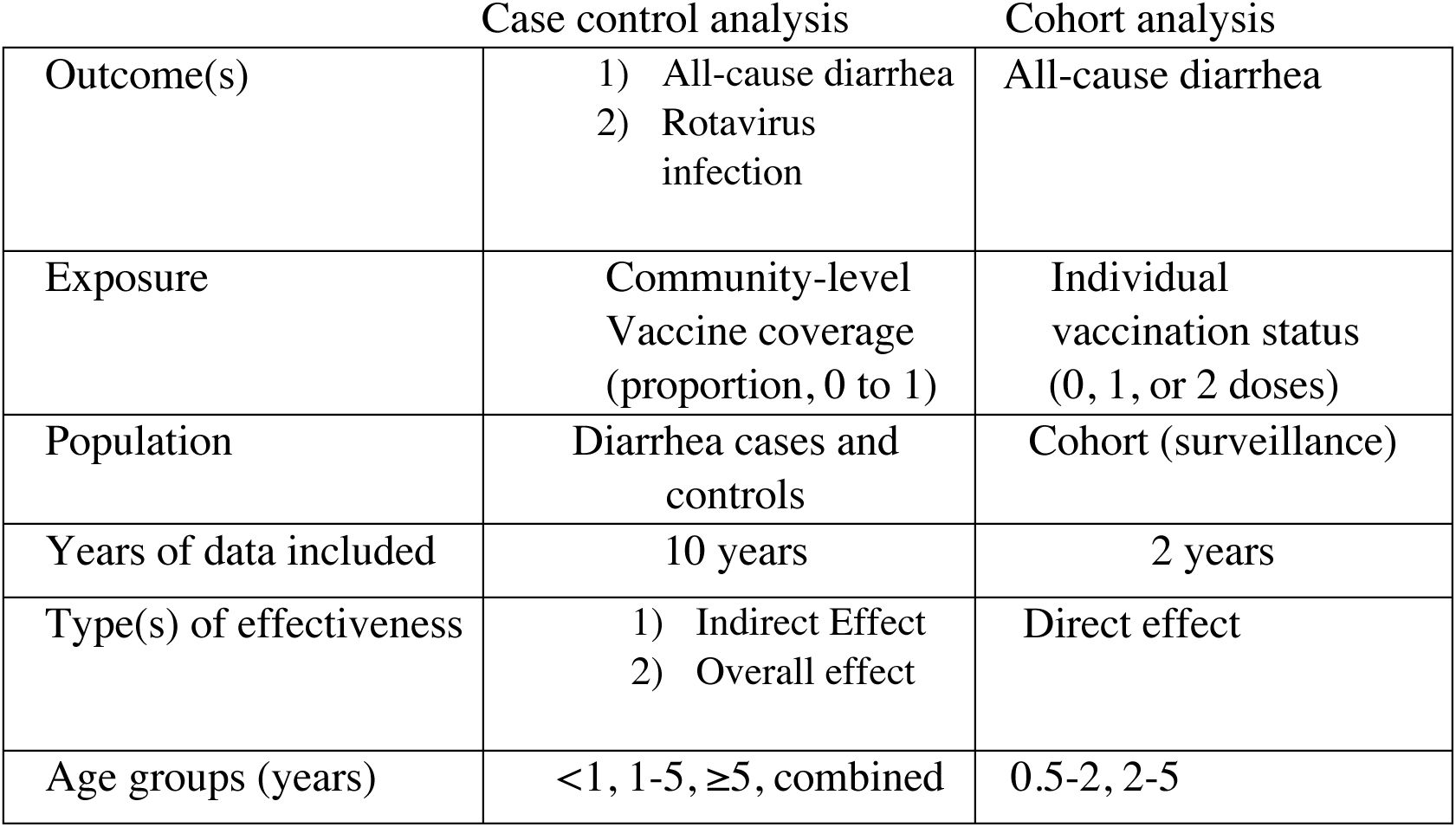
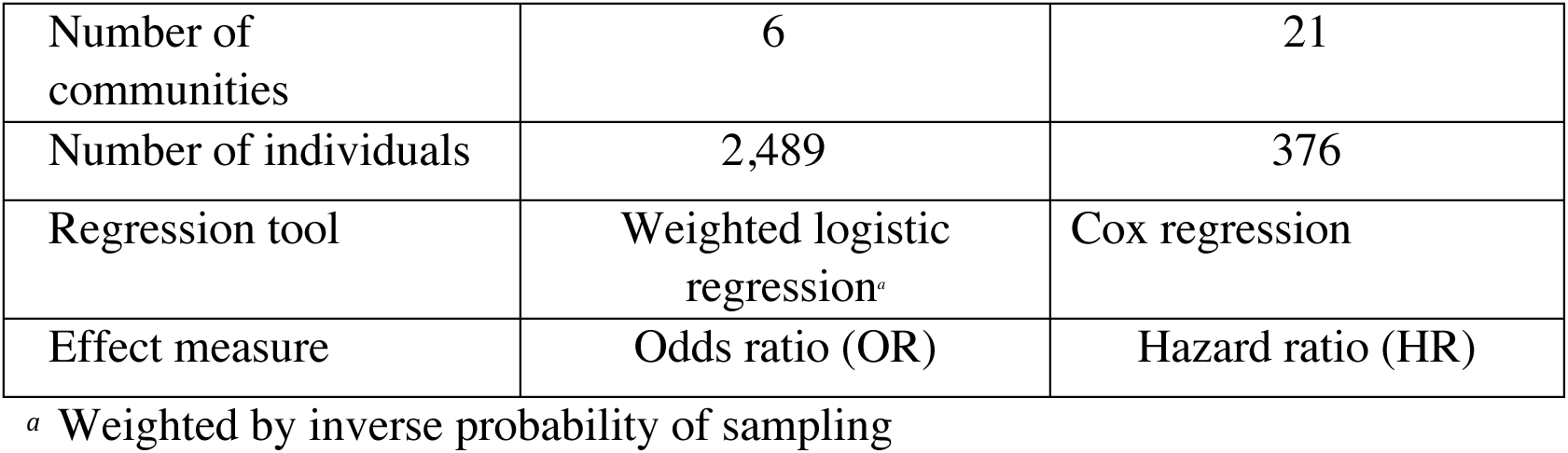
Summary of analytic approach. For the case-control analysis, only communities with at least one rotavirus infection in every cycle from the case control study were eligible for inclusion (total of 6 communities). For the cohort analysis, only communities with vaccine records were eligible for inclusion in the analysis (total of 21 communities). Selecting communities in this way would tend to bias our effect estimates towards the null. See supplement (section S3) for details.

|  | Case control analysis | Cohort analysis |
| --- | --- | --- |
| Outcome(s) | 1) All-cause diarrhea<br>2) Rotavirus infection | All-cause diarrhea |
| Exposure | Community-level Vaccine coverage (proportion, 0 to 1) | Individual vaccination status (0, 1, or 2 doses) |
| Population | Diarrhea cases and controls | Cohort (surveillance) |
| Years of data included | 10 years | 2 years |
| Type(s) of effectiveness | 1) Indirect Effect<br>2) Overall effect | Direct effect |
| Age groups (years) | <1, 1-5, $\geq 5$, combined | 0.5-2, 2-5 |
| Number of communities | 6 | 21 |
| Number of individuals | 2,489 | 376 |
| Regression tool | Weighted logistic regression<sup>a</sup> | Cox regression |
| Effect measure | Odds ratio (OR) | Hazard ratio (HR) |

<sup>a</sup> Weighted by inverse probability of sampling

^*a*^Weighted by inverse probability of sampling

For the case-control analysis, we estimated the association between vaccine coverage of two doses of Rotarix (estimated using the cohort data) on both all-cause diarrhea and rotavirus infection for the population overall as well as for three age groups separately: <1, 1-5, and ≥5. Children began to receive their second dose of vaccine during cycle 8 of data collection. Since we averaged coverage at the cycle level and each cycle lasted 9 calendar months on average, we lagged vaccine coverage by one cycle, so that the exposure variable reflected children actual vaccine coverage a few months prior to the start of each cycle rather than reflecting doses that were administered throughout the case control cycle. For the cohort analysis, we included children under five years who were age-eligible to be vaccinated. As a sensitivity analysis, we also considered the potential for chance confounding by seasonality to have biased the results. The effect estimates were extremely similar before and after adjustment for season so we did adjust our final models for season (see S5 for details).

For the case-control analysis, only communities that had at least one positive stool sample in each cycle were included (6 communities, 2,489 individuals). For the cohort analysis, only communities that had vaccine records were included. All statistical analyses were conducted in R (version 3.4). Weighted regression analyses used the package ‘survey,’ clustered logistic regression used the package ‘gee,’ and Cox regression analyses used the ‘OIsurv’ function from the package ‘stats’.

#### Case-control analysis

We compared risk of rotavirus infection and all-cause diarrhea over time and by age group to assess whether the vaccine was associated with a change in transmission patterns in our study region using data from six communities (n=2489, 150 rotavirus positive). All-cause diarrhea cases were defined as cases in the case-control study. Symptomatic rotavirus infections were defined as cases of diarrhea in the case-control study who tested positive for rotavirus. Asymptomatic rotavirus infections were defined as controls without diarrhea symptoms who tested positive for rotavirus.

We used individual-level logistic regression models, weighted by inverse probability of sampling, to compare the odds of rotavirus infection over time, using community coverage of two doses of Rotarix as our exposure. We conceptualized our data for this analysis as repeated cross-sectional surveys that provided an unbiased estimate of rotavirus infection and all-cause diarrhea within the community at each time point. We conducted this analysis separately by age group (<1 year, 1-5 years, and ≥5 years) and for all age groups combined. We calculated overall percent reduction in all-cause diarrhea and rotavirus infection for all age groups separately and the population overall (*Reduction* = (1 − *OR*) × 100).

#### Cohort analysis

We created a dynamic cohort of children enrolling all children with vaccine records born between August 2008 and September 2013, with children entering the cohort at six months of age (the latest age Rotarix was administered according to local policy). However, children born between 2008 and 2011 were older upon cohort entry (although they were still vaccinated before 6 months of age) because we did not have surveillance data before 2011 (see Figure 1). Based on census records, 819 children born in the 21 study communities between 2008 and 2013 were included in the analysis with surveillance data. Vaccine records and covariate data were available for 376 of these children (46%). We compared the time to first all-cause diarrhea case by vaccination status (2 doses, 1 dose, reference group=0 doses) for these 376 children using Cox regression to estimate the direct effect of vaccination (using the hazard ratio, HR).

Because vaccine records were not available for about half of our study population, we also compared children with and without vaccine records to assess whether these groups were comparable. While we focus on results for the case-control study assuming that coverage of rotavirus vaccination in the population overall was similar for children with and without records, we also considered alternative assumptions in sensitivity analyses (see S6). In general, the association between vaccine coverage on rotavirus infection diarrhea was similar but slightly stronger when vaccine coverage was lower among children without vaccine records than among children with records. We retained the assumption of similar coverage so that our results would be conservative.

#### Comparing Analysis Parts

Because both analyses used different communities, and some had substantial missing data, we ran Poisson models for the cohort analysis that used the count of all-cause diarrhea episodes as the outcome. We then compared the rate ratio from this model to the OR quantifying the association between vaccine coverage and all-cause diarrhea among children from the case-control analysis. For this comparison, we did not lag vaccine coverage so that the exposure windows would be comparable. Because the cohort analysis used only children with vaccine records from 21 communities whereas the case-control analysis used all cases and controls from 6 communities, comparing the results provides an internal consistency check (see S9 for details).

## Results

### Descriptive statistics

#### Case control data

*Vaccine coverage (exposure).* The average overall vaccine coverage was 74.6% for all cycles and was relatively constant over time (see supplement). At the community level, coverage estimates varied, ranging from 40·0% to 100%, with higher vaccine coverage in less remote villages.

*Time trends in all-cause diarrhea and rotavirus infection (outcomes).* While older children and adults have the lowest per-capita risk of rotavirus infection (Figure 2A), they explain a substantial proportion of the symptomatic infections due to their higher proportion in the population (Figure 2B). In this older age group, rotavirus was also a causative diarrheal pathogen (see Table S9). The total prevalence of rotavirus infection and all-cause diarrhea decreased after the vaccine was introduced, with young children <1 year of age showing strong reductions beginning in cycle 8 when the vaccine was first introduced and older age groups showing this benefit 1 cycle later (Figure 2A). Graphs of rotavirus infection (symptomatic and asymptomatic) and all-cause diarrhea over time are shown in Figure S-3.

**Figure 2.**
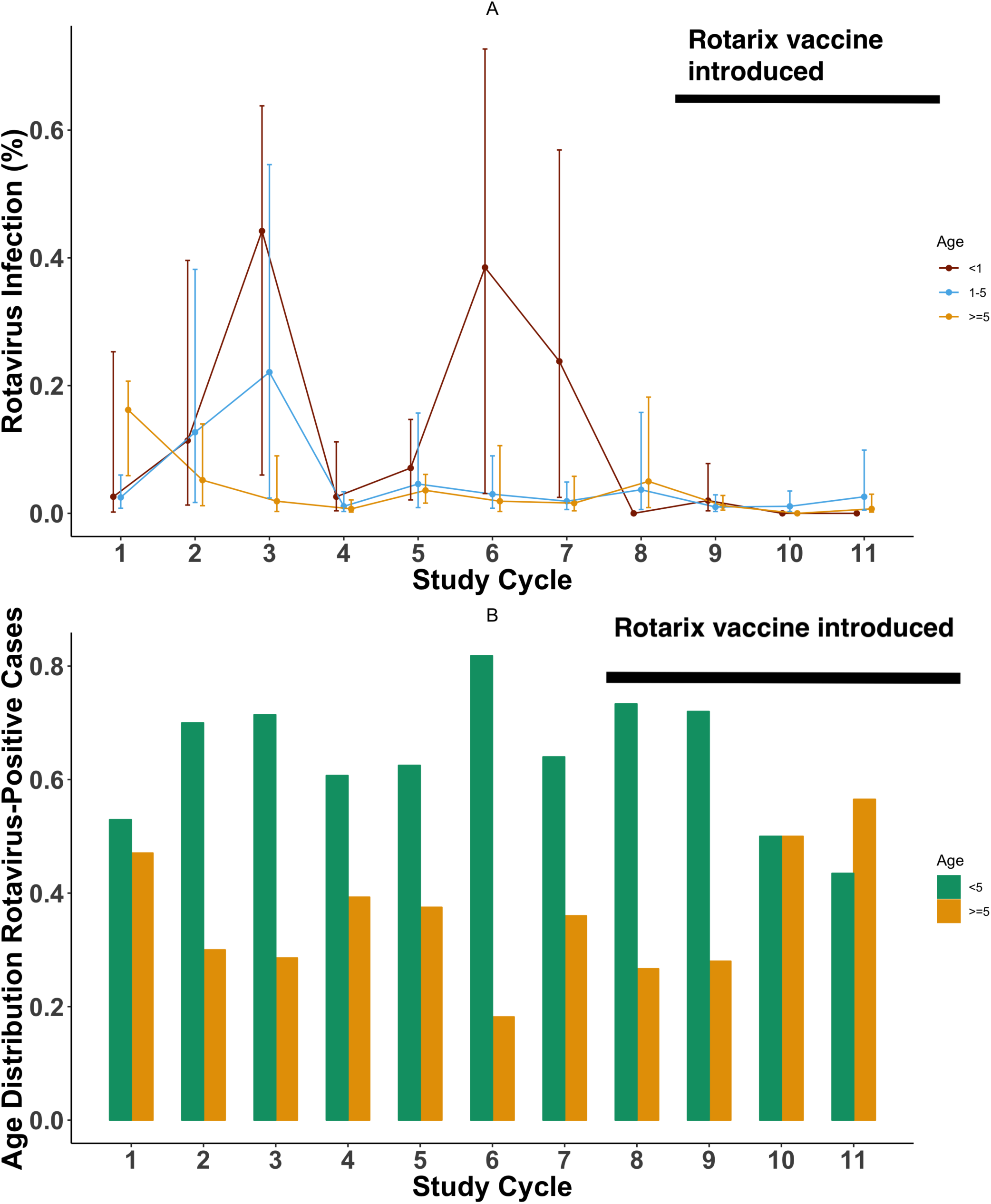
A) Percent of population infected with rotavirus by age group, B) Age distribution of symptomatic rotavirus positive individuals by study year. For panel A, children <1 year of age are shown in red, children between 1 and 5 years of age are shown in light blue, and older children (≥5 years) and adults are shown in gold. For panel B, older children and adults are shown in gold and children under 5 years of age are shown in dark green. The vaccine was first introduced in this region during cycle 8 (late in 2008).

*Covariates.* In the case-control study, the prevalence of rotavirus infection in the general population declined after vaccine introduction (4.9% to 0.8% in the full population, qualitative comparison, see Table 2). Over time, the prevalence of all-cause diarrhea also declined (2.2% to 1.6%) and the socioeconomic status improved, as indicated by smaller household size and higher average household education.

**Table 2.** Characteristics of study communities before (cycles 1-8) and after (cycles 9-11) the vaccine was introduced. All statistics are weighted by inverse probability of sampling and the total sample size (N) is the total population size of the six communities included in the analysis. Continuous variables are shown as mean (standard error) and categorical variables are shown as % (n).

|  | <u>Case-control analysis</u> |  |
| --- | --- | --- |
|  | Pre-vaccine<br>(N=20,660) | Post-vaccine<br>(N=7,497) |
| Symptomatic for diarrhea (%) | 2.2(446) | 1.6(122) |
| Male (%) | 51.3 (10,617) | 52.0 (3902) |
| Age (%) |  |  |
| <1 | 3.5(713) | 3.8(286) |
| 1-5 | 10.9(2260) | 10.4(777) |
| >5 | 84.9(17,548) | 85.6(6432) |
| Household size (n) | 6.75(2.89) | 6.34(2.83) |
| Highest household education (yrs) | 7.90(3.38) | 8.3(3.35) |
| Remoteness (%) |  |  |
| Close | 66.1(13,653) | 52.6(3944) |
| Medium | 5.6(1158) | 5.2(390) |
| Far | 28.3(5850) | 42.2(3164) |
| Rotavirus infection (%) | 4.9(845) | 0.8(60) |

#### Cohort data

Differences between vaccinated and unvaccinated children are shown in Table 3. Children with vaccine records available had a higher hazard of illness than children without vaccine records and a higher proportion of them became sick during the surveillance period. While not significant, children without vaccine records also tended to come from houses with higher education (*p* = 0.064) and fewer children (p=0.079), both of which may indicate higher socioeconomic status.

**Table 3.** Surveillance data descriptive statistics. Continuous variables are shown as mean (standard deviation), categorical variables and risks are shown as %(n), and rates are shown as rates(number of events). A * represents a significant difference between children with and without vaccine records (comparing columns 1 and 2)

|  | No vaccine record<br>(N=443) | Total<br>(N=376) | Zero Doses<br>(N=47) | One Dose<br>(N=58) | Two Doses<br>(N=271) |
| --- | --- | --- | --- | --- | --- |
| Remoteness (%) |  |  |  |  |  |
| Close | 46.7 (207) | 48.7 (183) | 59.6(28) | 27.6(16) | 51.3(139) |
| Medium | 6.6 (29) | 8.5 (32) | 10.6(5) | 6.9(4) | 8.5(23) |
| Far | 46.7 (207) | 42.8 (161) | 29.8(14) | 65.5(38) | 40.2(109) |
| Age (yrs) |  |  |  |  |  |
| Cohort Entry | 1.49(1.00) | 1.41(0.92) | 1.66(1.01) | 1.58(1.05) | 1.33(0.86) |

Table 3. Surveillance data descriptive statistics.
|  |  |  |  |  |  |
| --- | --- | --- | --- | --- | --- |
| First Diarrheal Case | 1.89(1.04) | 1.79(0.96) | 1.92(0.87) | 1.61(0.86) | 1.80(1.00) |
| BCG Vaccination (%) | N/A | 94.1(353) | 74.5(35) | 94.8(55) | 98.5(266) |
| Male (%) | 48.7(182) | 47.6(156) | 63.2(24) | 43.2(19) | 45.9(113) |
| Household Size (total N) | 6.36(3.28) | 6.61(3.57) | 6.89(4.51) | 6.93(3.38) | 6.51(3.45) |
| Household Size (children) | 3.16(1.75) | 3.41(1.89) | 3.31(2.29) | 3.68(1.81) | 3.38(1.85) |
| Highest Household Education (yrs) | 8.79(3.56) | 8.29(3.27) | 8.56(2.81) | 8.26(3.60) | 8.26(3.28) |
| Overall Illness Rate (per 1,000 person-weeks) |  |  |  |  |  |
| Diarrhea | 14.28 (246) | 18.0(271)* | 17.8(36) | 19.9(49) | 17.6(186) |
| Non-diarrhea | 3.7(61) | 5.71(86)* | 4.94(10) | 8.10(20) | 5.29(56) |
| Any | 18.4(307) | 23.7(357)* | 22.7(46) | 28.0(69) | 22.9(242) |
| Person-weeks | 16,672 | 15,071 | 2025 | 2468 | 10,578 |
| Overall illness risk (%) |  |  |  |  |  |
| Diarrhea | 27.4(121) | 36.3(137)* | 38.3(18) | 37.9(22) | 35.7(92) |
| Non-diarrhea | 6.1(27) | 8.8(33) | 6.4(3) | 13.8(8) | 8.1(22) |
| Any | 31.4(139) | 41.1(155)* | 38.3(18) | 43.1(25) | 41.2(112) |
| Length diarrhea episode (days) | 2.61(2.02) | 2.47(1.98) | 2.13(1.95) | 2.26(1.51) | 2.61(2.05) |
| Length first diarrhea episode (days) | 3.40(1.84) | 3.20(1.79) | 2.74(1.63) | 3.18(1.82) | 3.29(1.82) |

While some children may not have vaccine records because they were not vaccinated, some of these vaccine records may have been lost and many may have been vaccinated elsewhere, particularly Borbón, the nearest city in the region and location of the region’s primary health center. Given that travel patterns in our study region are positively associated with socioeconomic status among adults [19], parents of children without vaccine records, who had higher education levels, may have brought their children to Borbón to receive vaccines. While not all children had dates associated with their vaccine records, children generally received both doses by six months of age (see S.1 and S.3). For communities included in the case-control analysis, we had vaccine coverage data for 43.9% of eligible children in cycle 8, 57.6% of eligible children in cycle 9, and 78.6% of eligible children in cycle 10 (used to define exposure for cycles 9, 10, and 11 respectively; see S-2 for vaccine availability by community).

### Case control analysis

Community-level vaccine coverage was inversely associated with rotavirus infection, with individuals living in communities with 100% vaccine coverage having 0.145 times the odds of rotavirus infection compared with individuals living in communities with 0% coverage (Table 4), corresponding to a reduction of 85.5% (95% CI: 61.1%, 94.5%) (Table 5). The reduction in asymptomatic infection comparing exposure to 100% vaccine coverage with 0% coverage was 90.1% (95% CI: 56.9-97.7%) and 48.3% (95% CI: 0.03-73.1%) for symptomatic infection (Table 5).

**Table 4.**
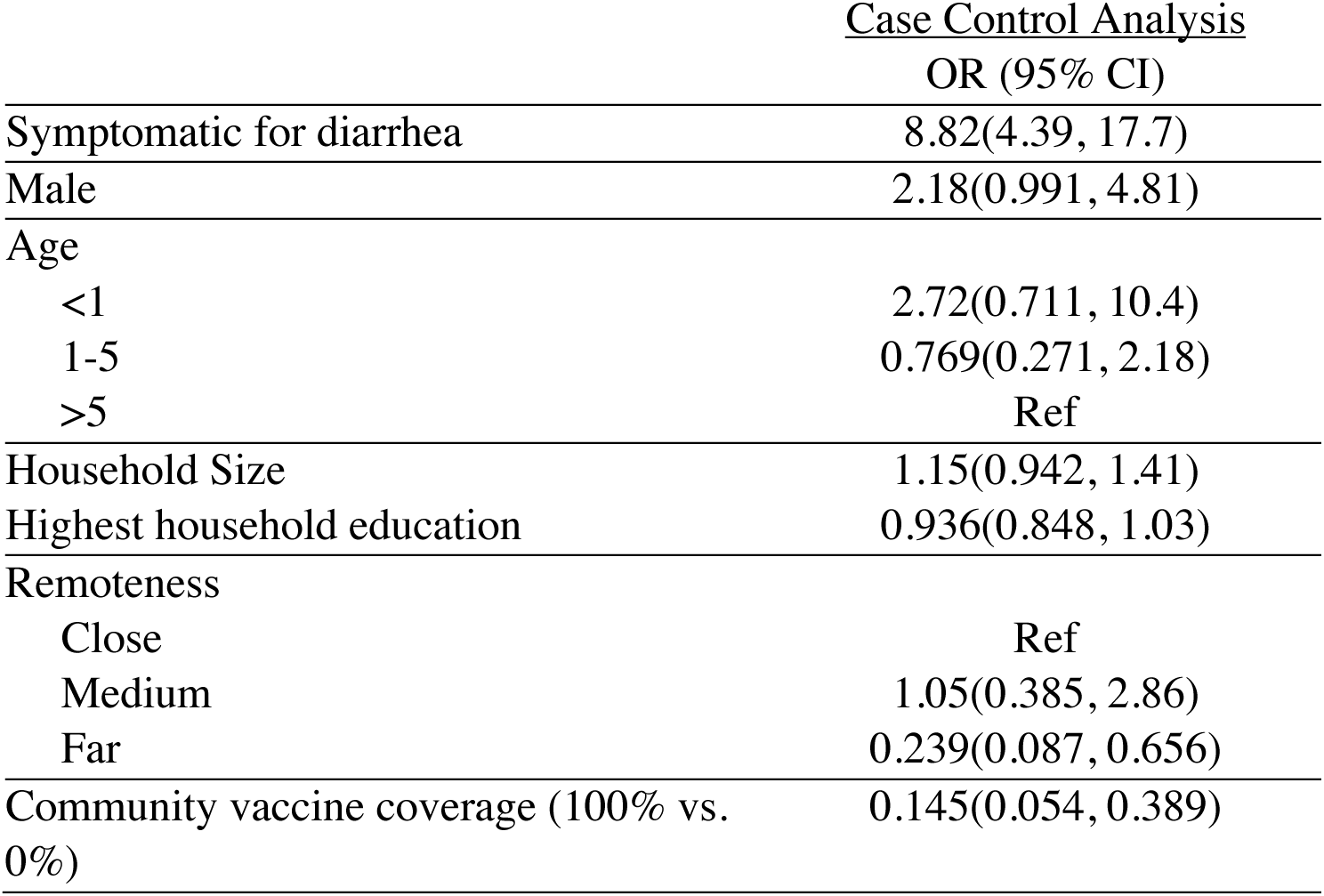
Logistic regression of rotavirus infection (combined symptomatic and asymptomatic). All cells represent OR (95% CI). Reference groups for categorical variables are noted above. OR for vaccine coverage compares 0% to 100%. Part 2 regression results are adjusted for whether or not infection was symptomatic, gender, age, household size, highest household education, and remoteness. Part 3 regression results are adjusted for all variables in part 2 except symptomatic (because all individuals in the part 3 analysis had symptomatic diarrhea).

| Case Control Analysis |  |
| --- | --- |
|  | OR (95% CI) |
| Symptomatic for diarrhea | 8.82(4.39, 17.7) |
| Male | 2.18(0.991, 4.81) |
| Age |  |
| <1 | 2.72(0.711, 10.4) |
| 1-5 | 0.769(0.271, 2.18) |
| >5 | Ref |
| Household Size | 1.15(0.942, 1.41) |
| Highest household education | 0.936(0.848, 1.03) |
| Remoteness |  |
| Close | Ref |
| Medium | 1.05(0.385, 2.86) |
| Far | 0.239(0.087, 0.656) |
| Community vaccine coverage (100% vs. 0%) | 0.145(0.054, 0.389) |

While the strongest effect was observed among young children (<1 year) (Table 5), all age groups had reduced risk of rotavirus infection, including people who were too old to have been vaccinated (indicating indirect effects).

**Table 5.** Overall effect of vaccination by age group, *VE* = (1 − *OR*) × 100. Models for rotavirus infection (columns 2-4) are adjusted for symptomatic/asymptomatic status, remoteness, gender, household size, and highest household education. Models for all-cause diarrhea (column 5) are adjusted for all variables in columns 2-4 except rotavirus infection. Models for the total population (row 4) are adjusted for age.

| Age group | Rotavirus Infection Risk |  |  | All-cause diarrhea |
| --- | --- | --- | --- | --- |
|  | Any | Symptomatic | Asymptomatic |  |
| <1 | 98.1% (-8.5%, 100.0%) | 77.5% (-43.4%, 96.5%) | 100% (N/A)* | 69.8% (-11.1%, 91.8%) |
| 1-5 | 82.8% (31.8%, 95.7%) | 34.1% (-81.6%, 76.1%) | 100% (N/A)* | 23.9% (-37.0%, 57.7%) |
| ≥5 | 84.5% (48.2%, 95.4%) | 49.2% (-28.5%, 79.9%) | 86.0% (40.7%, 96.7%) | 2.0% (-52.4%, 36.9%) |
| Total population | 85.5% (61.1%, 94.6%) | 48.3% (.03%, 73.1%) | 90.1% (56.9%, 97.7%) | 22.8% (-11.8%, 46.6%) |

### Cohort analysis

Among young children (6 months-2 years), rotavirus vaccination was associated with a decreased hazard of all-cause diarrhea (*HR*=0.429, 95% CI: 0.221, 0.834; Figure 3 and Table S6). Among older children born closer to the time of vaccine introduction (2-5 years), two doses of vaccine was also associated with a decreased hazard, but this association did not reach significance (*HR*=0.565, 95% CI: 0.121, 2.63). While children who received one dose of rotavirus also tended to have a decreased hazard of diarrhea compared to unvaccinated children during the first two years of life, this association was not significant (*HR*=0.577, 95% CI: 0.256, 1.30), which likely reflects the small sample size of this group.

**Figure 3.**
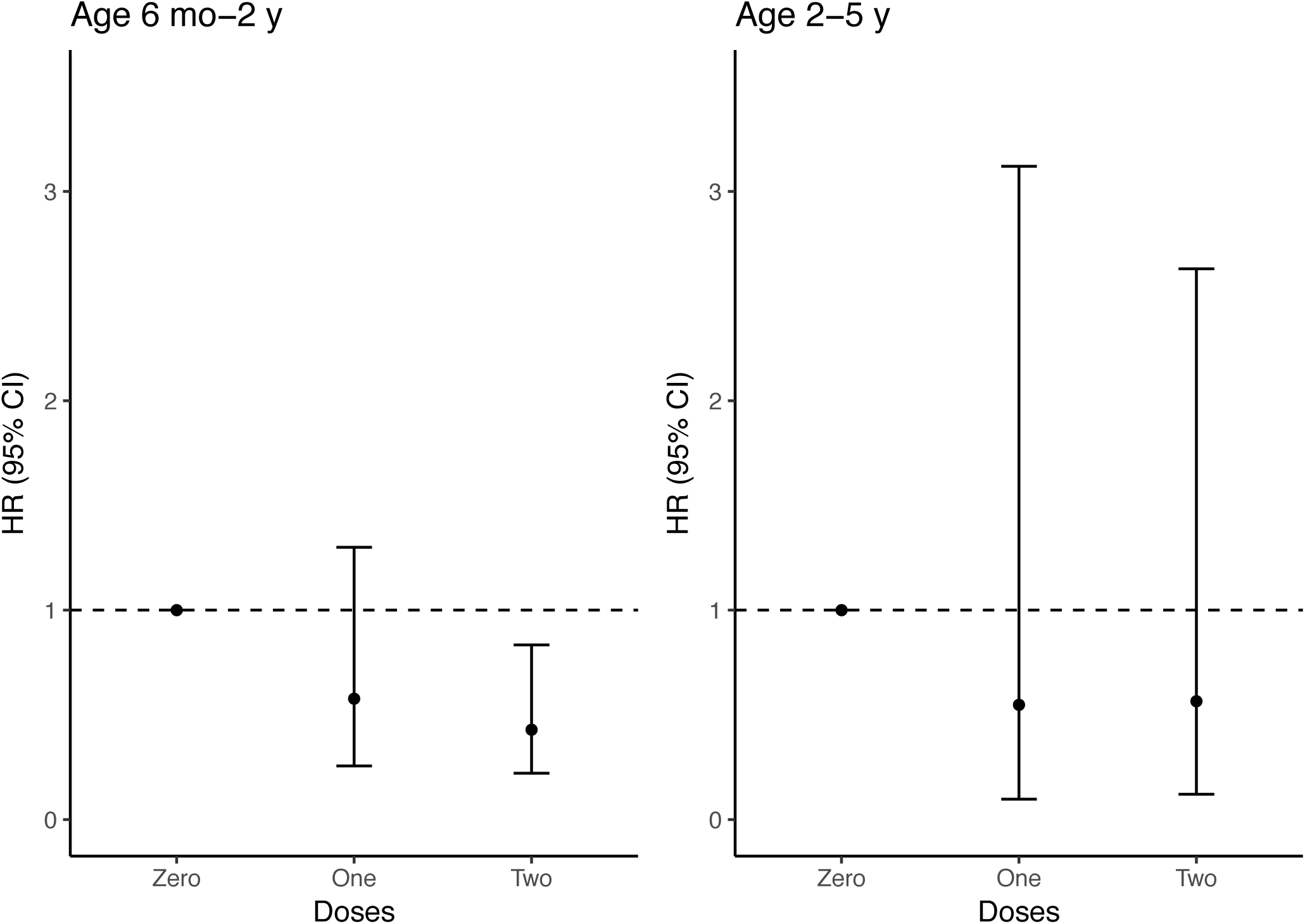
Direct effect of rotavirus vaccination on the rate all-cause diarrhea by age, calculated using Cox regression. Hazard ratios (HR) are adjusted for community remoteness, gender, highest household education, number of children in the household, and BCG vaccination. For children under two years, the reference group is children aged 6 months-2 years who received zero doses of vaccine. For children aged 2-5 years, the reference group is children aged 2-5 years who received zero doses of vaccine.

### Comparing analysis parts

Comparing the impact of vaccination on the rate of all-cause diarrhea, the point estimates estimated in both analyses were highly similar (See table S-9, among children <2 years, 39.8% reduction for the rate of all-cause diarrhea from the cohort compared with 46.3% for the case-control study).

## Discussion

In this study, we have shown that Rotarix has strong effectiveness on both rotavirus infection and all-cause diarrhea in a rural, nonclinical population in Ecuador. These effects are strongest among young children who are generally at highest risk of severe disease. However, we also found indirect effects on both outcomes among older children and adults who could not be vaccinated, indicating indirect effects. This is most likely due to suppression of overall transmission among all age groups.

Specifically, children < 1 year old tended to have reduced illness (77.5% reduction in symptomatic rotavirus infection and 69.8% reduction for all-cause diarrhea; Table 5). This effect size for symptomatic infection is in close agreement with that reported in randomized clinical trials in Latin America (VE 81%) [9]. While we estimate higher direct effectiveness for all-cause diarrhea (57.1%) than the 39-42% efficacy reported previously against hospitalization for all-cause diarrhea [9, 24], our confidence intervals overlap the estimates provided in these prior studies. The smaller and nonsignificant effect size for older children in the cohort is likely partially attributable to naturally-acquired immunity during the years between vaccination and cohort entry [25].

All age groups were significantly protected from asymptomatic infection, including the oldest, unvaccinated age group (85.5% reduction in rotavirus infection for 100% versus 0% coverage). This result implies that Rotarix vaccination suppresses not only severe disease but also overall transmission and helps explain why we and others have observed indirect effects of vaccination in older age groups [16–18]. We find that this benefit is attainable in low resource settings. This suppression of overall infection also suggests that prior studies, which focused on severe rotavirus infection in a clinical setting, may have underestimated the total impact of vaccination on rotavirus infection.

## Strengths and Limitations

Due to our retrospective study design, we had high levels of missing data for the cohort analysis decreasing our sample size, which could have also biased our estimates for vaccine coverage for the communities overall. However, the total effect of coverage on vaccination were robust to a variety of different coverage assumptions. When coverage was lower among children without vaccine records compared to those with records, our results were slightly stronger and the association between vaccine coverage and rotavirus infection became significant for young children. Moreover, given that the population level effects of vaccination were strongly dependent on our vaccine coverage estimates for all age groups rather than vaccine introduction alone (see S.3) and that parts 1 and 2 found similar results, it is likely that the actual vaccine coverage within our study communities is at least proportional to the coverage estimates from children with vaccine records.

As with any non-randomized study, our results are also potentially subject to bias from secular trends and unmeasured confounding. We have previously shown many community characteristics changed over this time period, including socio-economic status and mobility/migration, with impacts on rotavirus infection over the same period [19]. However, any confounder would need to be proportional to community vaccine coverage to explain the observed associations.

Because the case-control study was focused on symptomatic diarrhea, our confidence intervals for asymptomatic infection are wide and we cannot determine with certainty relative effectiveness on asymptomatic versus symptomatic infection. While uncertain, the higher point estimate against asymptomatic infection in this study may result from dose-dependent effects of vaccination, where the vaccine has higher protection against lower dose exposures. Additionally, the method used to detect rotavirus infections is somewhat less sensitive than RT-PCR and may have failed to detect asymptomatic rotavirus infections with lower shedding rates [26]. For this reason, our overall statistical power may have been reduced for asymptomatic infections in general and we may also have overestimated the effect of vaccination on asymptomatic infections with lower shedding rates. However, the asymptomatic infections with sufficient shedding rates to be detected by our methodology are probably those that are most relevant for secondary transmission. Despite these limitations, our results strongly suggest that Rotarix vaccination is protective against milder rotavirus infections, including asymptomatic infection, in addition to its effects on severe rotavirus. At a minimum, our results suggest that vaccination reduces shedding to below to the detection limit for the EIA assay, which would be expected to dampen overall transmission.

## Conclusions

Rotarix vaccination has substantially reduced both rotavirus infection and all-cause diarrhea in a rural region of Ecuador, despite significant challenges to vaccination. Much of this effect is driven by suppression of overall rotavirus transmission at a population level, including among older age groups who were too old to be vaccinated. The impact of rotavirus vaccination may be higher than previously estimated given its effect on milder diarrheal disease and asymptomatic infections.

## Supporting information

## Acknowledgements

We would like to thank the Ministry of Health for providing vaccine data and the EcoDess field team for their valuable contribution collecting the data.

