## Supplementary material for "Effectiveness of live attenuated monovalent human rotavirus vaccination in rural Ecuador, 2008-2013"

November 29, 2018

#### S1 Vaccine Data Collection Details

We collected vaccine data from a local health system. Vaccine records were available for 21 communities. The fraction of community members with vaccine records is shown in table S-1 and the coverage of two doses of vaccine among children with records by community is shown in table S-2. For both tables, communities that were included in the case-control analysis are shown in bold. In general, coverage was highest in close villages (see Table S-3). Most children had completed their vaccine dosing by six months of age. Of children who ultimately received only one dose, almost all had received this dose by six months of age and most children who received both doses of vaccine had received both doses by 6 months of age (Table S-4). Based on this information, we calculated vaccine coverage among children with records using the number of children with vaccine records who were six months old in a given case-control study cycle as the denominator and the numerator was the number of age-eligible children who actually received their second dose of vaccine. While the number of children with vaccine records in each study community was small, this constituted a substantial fraction of the population. For the communities included in the case-control analysis, the availability of vaccine records was higher on average, with average record availability of 44%, 58%, 79% and 69% of communities included in the analysis for cycles 8, 9, 10, and 11 respectively (community 2 is not included in the record availability calculation because we did not include cycles in which the number of vaccine records was 0 from the regression analysis).

#### S2 Defining vaccine effectiveness parameters

We use the counterfactual framework described by Halloran and Hudgens [1] to estimate vaccine effectiveness. For all equations, we let two doses of vaccine be defined by a parameter called $vaccinated$, which takes a value of 1 for fully vaccinated individuals and a value of 0 for unvaccinated individuals. The variables $\alpha$ and $\alpha'$ define the regional or community coverage of vaccination. Each variable is defined in its corresponding section. In all equations, $\bar{y}$ is a statistic that quantifies the average outcome (rate or odds depending on the model) for a given covariate pattern. All effectiveness parameters are conceptually defined based on a counterfactual contrast, but all variables in these equations are represented by statistics within our sample because all individuals at risk are not observed (i.e., our statistics are random variables that depend on our sampling process). Analogously, the vaccine effectiveness quantities are represented with an overhead bar

| Community | Cycle 8 | Cycle 9 | Cycle 10 | Cycle 11 |
|---|---|---|---|---|
| Close Communities | | | | |
| **1** | **58.3 (7/12)** | **45.0 (9/20)** | **44.4 (8/18)** | **81.8 (9/11)** |
| **2** | **43.8 (7/16)** | **61.9 (13/21)** | **34.3 (12/35)** | **0 (0/22)** |
| **3** | **44.4 (12/27)** | **67.8 (25/37)** | **80.0 (28/35)** | **85.0 (17/20)** |
| 4 | 0 (0/7) | 45.5 (5/11) | 50.0 (8/16) | 0 (0/14) |
| 5 | 30.0 (3/10) | 40.0 (4/10) | 44.4 (8/18) | 0 (0/12) |
| *Total* | 40.2 (29/72) | 56.6 (56/99) | 52.5 (64/122) | 32.9(26/79) |
| Medium Communities | | | | |
| 6 | 50.0 (1/2) | 100 (2/2) | 50.0 (2/4) | 100 (1/1) |
| **7** | **66.7 (2/3)** | **54.5 (6/11)** | **100 (6/6)** | **60.0 (3/5)** |
| 8 | 100 (1/1) | 100 (3/3) | 40.0 (2/5) | 0 (0/7) |
| 9 | 50.0 (1/2) | 80.0 (4/5) | 66.7 (2/3) | 50.0 (1/2) |
| 10 | 100 (1/1) | 66.7 (2/3) | 50.0 (2/4) | 100 (1/1) |
| *Total* | 66.7 (6/9) | 70.8 (17/24) | 63.6 (14/22) | 37.5 (6/16) |
| Far Communities | | | | |
| **11** | **19 (3/16)** | **36.4 (4/11)** | **29.4 (5/17)** | **50.0 (3/6)** |
| 12 | 100 (7/7) | 100 (6/6) | 66.7 (2/3) | 100 (4/4) |
| 13 | 0 (0/1) | 33.3 (1/3) | 75.0 (3/4) | 33.3 (1/3) |
| **14** | **62.5 (5/8)** | **60.0 (15/25)** | **45.0 (9/20)** | **46.2 (6/13)** |
| 15 | 75.0 (3/4) | 0 (0/3) | 0 (0/3) | 0 (0/8) |
| 16 | 0 (0/3) | 12.5 (1/8) | 40.0 (4/10) | 71.4 (5/7) |
| 17 | 76.9 (10/13) | 66.7 (12/18) | 50.0 (6/12) | 75.0 (12/16) |
| 18 | 0 (0/2) | 40.0 (2/5) | 50.0 (2/4) | 60.0 (3/5) |
| 19 | 60.0 (3/5) | 50.0 (3/6) | 25.0 (5/20) | 46.2 (6/13) |
| 20 | 0 (0/4) | 0 (0/3) | 0 (0/7) | 36.4 (4/11) |
| 21 | 0 (0/12) | 18.2 (2/11) | 30.0 (6/20) | 72.7 (8/11) |
| *Total* | 41.3 (31/75) | 47.4 (46/97) | 35.0 (42/120) | 53.6(52/97) |

Table S-1: Vaccine record availability by community. Each cell is presented as Percent with Vaccine Records (Number with Vaccine Records/Number Eligible). Communities that were included in the rotavirus positivity analysis are shown in bold

| Community | Cycle 8 | Cycle 9 | Cycle 10 | Cycle 11 |
|---|---|---|---|---|
| Close Communities | | | | |
| **1** | **85.7 (6/7)** | **77.8 (7/9)** | **87.5 (7/8)** | **88.9 (8/9)** |
| **2** | **57.1 (4/7)** | **46.2 (6/13)** | **50.0 (6/12)** | **N/A** |
| **3** | **100 (12/12)** | **88.0 (22/25)** | **100.0 (28/28)** | **100.0 (17/17)** |
| 4 | N/A | 20.0 (1/5) | 37.5 (3/8) | N/A |
| 5 | 100 (3/3) | 25.0 (1/4) | 37.5 (3/8) | N/A |
| *Total* | 86.2 (25/29) | 66.1 (37/56) | 73.4 (47/64) | 96.2 (25/26) |
| Medium Communities | | | | |
| 6 | 0 (0/1) | 100 (2/2) | 100 (2/2) | 100 (1/1) |
| **7** | **50.0 (1/2)** | **33.3 (2/6)** | **100 (6/6)** | **100 (3/3)** |
| 8 | 0 (0/1) | 66.7 (2/3) | 50.0 (1/2) | N/A |
| 9 | 100 (1/1) | 75.0 (3/4) | 50.0 (1/2) | 100 (1/1) |
| 10 | 100 (1/1) | 100 (2/2) | 100 (2/2) | 100 (1/1) |
| *Total* | 50.0 (3/6) | 64.7 (11/17) | 85.7 (12/14) | 100 (6/6) |
| Far Communities | | | | |
| **11** | **100 (3/3)** | **100 (4/4)** | **100 (5/5)** | **100 (3/3)** |
| 12 | 57.1 (4/7) | 50.0 (3/6) | 50.0 (1/2) | 75.0 (3/4) |
| 13 | N/A | 100 (1/1) | 66.7 (2/3) | 100 (1/1) |
| **14** | **100 (5/5)** | **73.3 (11/15)** | **88.9 (8/9)** | **83.3 (5/6)** |
| 15 | 66.7 (2/3) | N/A | N/A | N/A |
| 16 | N/A | 100 (1/1) | 100 (4/4) | 60.0 (3/5) |
| 17 | 70.0 (7/10) | 66.7 (8/12) | 66.7 (4/6) | 100 (12/12) |
| 18 | N/A | 50.0 (1/2) | 100 (2/2) | 33.3 (1/3) |
| 19 | 100 (3/3) | 66.7 (2/3) | 60.0 (3/5) | 33.3 (2/6) |
| 20 | N/A | N/A | N/A | 25.0 (1/4) |
| 21 | N/A | 0 (0/2) | 33.3 (2/6) | 50.0 (4/8) |
| *Total* | 77.4 (24/31) | 67.4 (31/46) | 82.1 (32/39) | 67.3 (35/52) |

Table S-2: Coverage of two doses of rotavirus vaccine among children with vaccine records. All cells show % Vaccinated (n vaccinated/n with vaccine records). A value of 'N/A' indicates that no vaccine records were available for that cycle.

| Remoteness | Cycle 8 | Cycle 9 | Cycle 10 | Cycle 11 |
|---|---|---|---|---|
| Close Communities | | | | |
| Fraction with Records | 40.2% (29/72) | 56.6% (56/99) | 52.5% (64/122) | 32.9% (26/79) |
| Coverage of two doses | 86.2% (25/29) | 66.1% (37/56) | 73.4% (47/64) | 96.2% (25/26) |
| Medium Communities | | | | |
| Fraction with Records | 66.7% (6/9) | 70.8% (17/24) | 63.6% (14/22) | 37.5% (6/16) |
| Coverage of two doses | 50.0% (3/6) | 64.7%(11/17) | 85.7% (12/14) | 100% (6/6) |
| Far Communities | | | | |
| Fraction with Records | 41.3% (31/75) | 47.4% (46/97) | 35.0 % (42/120) | 53.6% (52/97) |
| Coverage of two doses | 77.4% (24/31) | 67.4% (31/46) | 82.1% (32/39) | 67.3% (35/52) |
| Overall Coverage | 78.8% (52/66) | 66.4% (79/119) | 77.8% (91/117) | 78.6% (66/84) |

Table S-3: Vaccine coverage over time by community remoteness

| | Children ultimately receiving one dose (N=47) | Children ultimately receiving two doses (N=264) |
|---|---|---|
| Records with date information | 76.7% (n=36) | 73.8% (n=195) |
| Vaccinated by six months of age | 91.7% (n=43) | 88.7% (n=173) |
| Vaccinated by seven months of age | 100% (n=46) | 96.2% (n=188) |

Table S-4: Fraction of children receiving Rotarix vaccine by age. Column 1 is for children who were considered to have received one dose in the regression model and corresponds to the date by which they received their one and only dose and column 2 is for children who ultimately received two doses and corresponds to the completion date of their second dose of vaccine. These results are shown for all children for whom we had vaccine records, which includes some children who were not in the full regression analysis due to missing data on socioeconomic indicators.

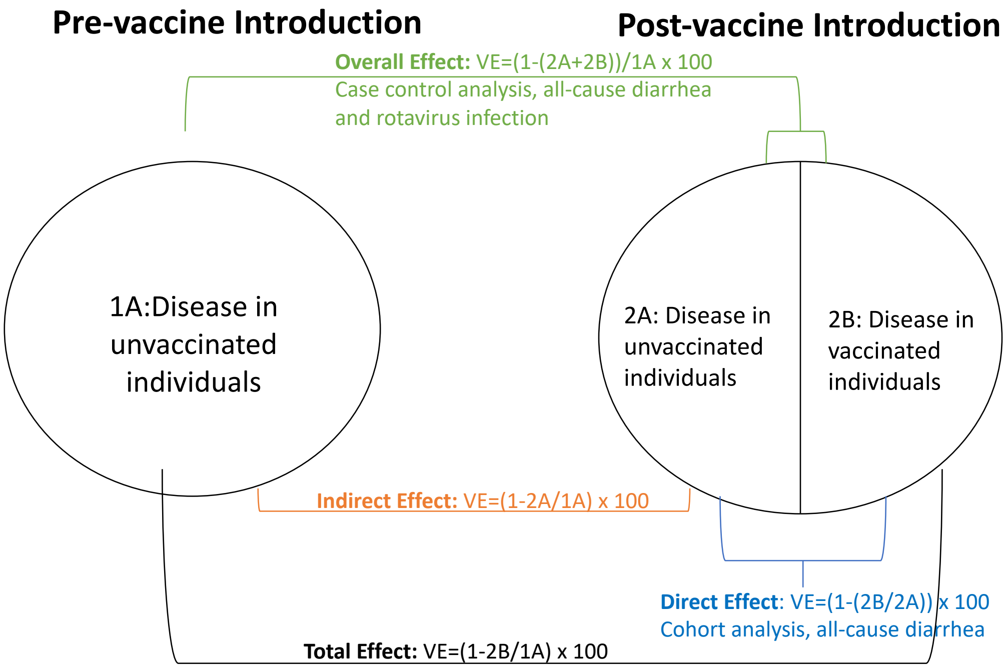


Figure S-1: Diagram showing how each vaccine effectiveness measure was calculated. This figure was adapted from Panozzo et al, 2014 [2]. In this analysis, we estimated the overall effect of vaccination using data from the case control study (green) and the direct effect using data from the diarrheal disease surveillance to conduct a cohort analysis (blue). While not directly estimated, the overall effect of vaccination is equivalent to the indirect effect (orange) for older children and adults (age $\geq 5$) because there were no vaccinated individuals in that age group (i.e., 2B=0). The total effect (black) was not directly estimated, but is approximated by the overall effect for younger age groups when vaccine coverage is high (i.e., 2A is approximately equal to 0).

to indicate that the dependence of the estimated effects on our sampling process.

Vaccine effectiveness measures can be classified into four groups: direct effects, indirect effects, overall effects, and total effects. A diagram of each of these possible effectiveness measures and how they were estimated in our study is shown in Figure S-1.

Because all doses of rotavirus vaccine that were used to estimate vaccine coverage were received gradually over the course of the study cycle, we did not expect for the indirect effect of a given level of coverage for the general population to be present until the following case control cycle. However, because young children who were vaccinated should be protected soon after receiving their vaccine, coverage might be protective in the cycle in which coverage was estimated for the youngest age group (age<1 year). To account for this tendency, we lagged coverage by one study cycle for all regression models for analysis parts 1 and 2. However, for age-stratified analyses for children <1 year of age, we present both lagged and un-lagged regression results.

#### S2.1 Direct Effect (DE)

The general equation for calculating direct effects is shown below.

$$\overline{DE}(\alpha) = \bar{y}(vaccinated = 0|\alpha) - \bar{y}(vaccinated = 1|\alpha)$$

*Under 5 years of age* In our analysis, we estimate the direct effect of vaccination on all cause diarrhea using Cox regression analysis. To estimate this quantity, we compare the time to first all-cause diarrhea case for fully vaccinated children with children who were not vaccinated. We estimate this quantity for all study communities simultaneously, so $\alpha$ is equal to the average vaccine coverage after introduction (around 75%). We estimate this quantity separately for 2 doses and 1 dose and code the model as below using dummy variables. We only estimate the effect of one dose for the direct effect–all other vaccine effectiveness measures at the population level are for coverage of two doses of vaccine. Each $\beta_{X_i}$ term is the multiplicative increase in the hazard of diarrhea for a one unit increase in covariate $X_i$ for continuous variables or an indicator variable for its presence/absence for binary variables.

$$\lambda(t) = \lambda_0(t)\exp(\beta_{2doses}X_{i,2doses} + \beta_{1dose}X_{i,1dose} + ...)$$

Because this equation estimates the average time to first all-cause diarrhea case for a given covariate pattern, the difference in outcomes is given by exponentiating $\beta_{2dose}$ for two doses of vaccine and $\beta_{1dose}$ for one dose of vaccine. When the interaction term with age>2 years is added to the model, this contrast gives the vaccine efficacy for children under 2 years of age. So the indirect effect of vaccination is given by $(1 - \exp(\beta_{2doses}) \times 100 = (1 - \mathrm{HR}_{2dose}) \times 100$ for two doses of vaccine and $(1 - \exp(\beta_{1dose}) \times 100 = (1 - \mathrm{HR}_{1dose}) \times 100$.

*Over 5 years of age* Because no one over the age of 5 is vaccinated with rotavirus, there is no direct effect of vaccination.

#### S2.2 Indirect Effect (IE)

The general equation for calculating indirect effects is shown below.

$$\overline{IE}(\alpha, \alpha') = \bar{y}(vaccinated = 0; \alpha) - \bar{y}(vaccinated = 0; \alpha')$$

*Under 5 years of age* Because we do not know the vaccination status of every child after the vaccine was introduced and because only 8 children who received 0 doses of vaccine were sampled by chance in the case control study after vaccine introduction, we are not able to estimate the indirect effect on this age group.

*Over 5 years of age* Because no one over the age of five is vaccinated with rotavirus, only the indirect effect of vaccination can be calculated for this age group and in this case the overall effect is equal to the indirect effect. In our analysis, we estimate this indirect using logistic regression analysis for the sub-sample of our case control population over 5 years of age.

We compare the effect of vaccine introduction in general, so that $\alpha$ takes a value of 0 and $\alpha'$ is equal to the vaccine coverage after introduction. To compare rotavirus infection and diarrheal illness before and after the vaccine was introduced, we can code a dummy variable for cycle which takes values of 0 before the vaccine was introduced and during the first cycle of introduction and takes a value equal to the coverage of two doses of vaccine among children in the prior cycle thereafter, coded as a proportion between 0 and 1. We lag this vaccine coverage variable to account for the fact that children must seroconvert before adults can be indirectly protected. For young children, we also considered an un-lagged version of vaccine coverage because children are likely to be protected soon after completing their vaccine series.

The logistic regression model for rotavirus infection and for diarrhea illness in the case control then has the following form:

$$\log\Big(\frac{P(Rota=1)}{1-P(Rota=1)}\Big) = \beta_0 + \beta_{coverage}X_{i,coverage} + ....$$

In this equation, the intercept term, $\beta_0$ is the estimated odds of rotavirus infection when all other covariates ($X_i$) are equal to 0. Then, each $\beta_{X_i}$ term is the multiplicative increase in the odds for a one unit increase in covariate $X_i$ for continuous variables or an indicator variable for it's presence/absence for binary variables. The variable $\varepsilon_i$ is the error not captured by our regression model.

As described in the method section, this regression analysis was weighted to account for complex sampling design. Because this equation estimates the average odds of rotavirus infection for a given covariate pattern, the difference in outcomes is given by exponentiating $\beta_{coverage}$ and the contrast is interpreted as the effect for 100% coverage of vaccine versus 0% coverage. So the indirect effect of vaccination is given by $(1 - \exp(\beta_{coverage})) \times 100 = (1 - OR) \times 100$. This equation takes the same form for all-cause diarrhea.

#### S2.3 Total Effect (TE)

The general equation for calculating total effects is shown below.

$$\overline{TE}(\alpha, \alpha') = \bar{y}(vaccinated = 0; \alpha) - \bar{y}(vaccinated = 1; \alpha')$$

*Under 5 years of age.* Unfortunately, we were not able to directly estimate this quantity with confidence using the data that we had.

Ideally, this quantity would be calculated by comparing the rates of diarrhea for children with two doses of vaccine after introduction with the general unvaccinated population before vaccination (i.e., comparing $\alpha =$

$0$ with $\alpha' = 0.76$). However, because we only had case control data for 53 children who were fully vaccinated and 3 children who were vaccinated with one dose of vaccine, we could not assess the effectiveness of vaccination on rotavirus infection.

To estimate this quantity for all cause diarrhea, We could have compared the rates of diarrhea in the 2011-2013 surveillance for fully vaccinated individuals with all children under five in the 2003-2007 surveillance but we elected not to do this because our missing data analysis suggested that children who did not have vaccine records were not comparable to those with vaccine records in 2011-2013. As such, in order to estimate this quantity with minimal bias, we would need to compare this two dose group with children who received as least one vaccine from 2003-2007. These earlier vaccine records had been lost, so we were not able to make this comparison. To deal with this issue, we could have matched children with vaccine data to other children during the surveillance period by education and community (accounting for the key differences between children with and without vaccine records in the 2011-2013 surveillance), but such an analysis would have required assuming that the two time periods were otherwise comparable (i.e., there were no secular trends not attributable to vaccination). We previously showed that the strains of rotavirus circulating in this region have changed rapidly over time and that strains varied widely between seasons in our case control analysis. For this reason, we were not comfortable making this assumption.

However, the total effect should be similar to the overall effect of vaccination for children less than 5 years of age when vaccine coverage is high (estimation procedure for overall effects is described below).

*Over 5 years of age.* Because individuals over five are not vaccinated, this quantity is not defined for this group.

#### S2.4 Overall Effect (OE)

The general equation for calculating overall effects is shown below.

$$\overline{OE}(\alpha, \alpha') = \bar{y}(\alpha) - \bar{y}(\alpha')$$

For all groups, we estimated the overall effects using case control data on rotavirus positivity. We considered both vaccine introduction (binary) and vaccine coverage (continuous with values between 0 and 1, but lagged by one cycle as described above) for each age group separately and for the population overall. We also considered an un-lagged regression model for the youngest age group, as described above.

We compared both the effect of vaccine introduction in general using a dummy variable as described above (so that $\alpha = 0$ and $\alpha' = 0.75$) and using the vaccine coverage among children with vaccine records, treating alpha as a continuous variable (so that $\alpha = 0$ and $\alpha' = 1$).

For vaccine introduction, this gives:

$$\log\left(\frac{P(Rota=1)}{1-P(Rota=1)}\right) = \beta_0 + \beta_{introduced} X_{i,introduced} + ....$$

For vaccine coverage, this gives:

$$\log\left(\frac{P(Rota=1)}{1-P(Rota=1)}\right) = \beta_0 + \beta_{coverage} X_{i,coverage} + ....$$

When vaccine coverage is high, the contrast of the $\beta_{coverage}$ term describes the comparison between 0 vaccine coverage and 100% vaccine coverage, in which case all eligible individuals are vaccinated so that the overall effect in this group approximates the total effect. We found that the vaccine coverage version of this variable better described the data and was also predictive of rotavirus infection in the cycles after the vaccine was introduced. For this reason, we used vaccine coverage as the exposure of interest for all regression models.

##### S2.5 Summary

A summary of the vaccine effectiveness measures calculated in this study and their sources is summarized in the table below.

#### S3 Supplemental information for case control analysis

##### S3.1 Population level trends in rotavirus infection and diarrhea over time

To calculate confidence intervals for population trends in infection and all-cause diarrhea, we fit intercept-only weighted logistic regression models separately for each outcome (4 outcomes: all-cause diarrhea, rotavirus infection, symptomatic rotavirus infection, and asymptomatic rotavirus infection, non-rotavirus diarrhea) and cycle (11 cycles) of the case control study for a total of 55 models. For rotavirus infection, we were also interested in prevalence by age group, so we subset this model by the three age groups (used in figure 2A) to produce a total of 33 models. Then, for each of these models, we used the standard logit transformation to produce confidence intervals for prevalence guaranteed to fall between 0 and 1. Representative equations for rotavirus infection are shown below:

$$\begin{aligned}
\bar{P}(\text{infection}) &= \frac{\exp(\beta_0)}{1+\exp(\beta_0)} \\
P_{LCL}(\text{infection}) &= \frac{\exp(\beta_0 - 1.96 \times SE(\beta_0))}{1+\exp(\beta_0 - 1.96 \times SE(\beta_0))} \\
P_{LCL}(\text{infection}) &= \frac{\exp(\beta_0 + 1.96 \times SE(\beta_0))}{1+\exp(\beta_0 + 1.96 \times SE(\beta_0))}
\end{aligned}$$

Table S-5: Source of effectiveness measures

| | Direct effect (DE) | Indirect Effect (IE) | Overall Effect (OE) |
|---|---|---|---|
| **Age $<$5** | | | |
| Rotavirus Infection | – | – | $(1 - \exp(\beta_{coverage})) \times 100$ ** |
| All-cause diarrhea | $(1 - \exp(\beta_{2dose})) \times 100$* | – | – |
| | $(1 - \exp(\beta_{1dose})) \times 100$ * | | |
| Non-diarrheal illness | $(1 - \exp(\beta_{2dose})) \times 100$* | – | – |
| | $(1 - \exp(\beta_{1dose})) \times 100$ * | | |
| **Age $\geq$5** | | | |
| Rotavirus Infection | N/A | $(1 - \exp(\beta_{introduced})) \times 100$** | Equal to IE |
| **General Population** | | | |
| Rotavirus Infection | N/A | – | $(1 - \exp(\beta_{coverage})) \times 100$** |

*Estimated using surveillance data and cox regression

** Estimated using case control data and weighted logistic regression

We found that the prevalence of diarrhea declined over time for both rotavirus and non-rotavirus diarrhea (Figure S-2). For rotavirus, this change was stronger for asymptomatic rotavirus infection than symptomatic rotavirus. However, the confidence intervals overlap and we were not adequately powered to determine if the effect on asymptomatic infection was stronger than on symptomatic infection (Figure S-3).

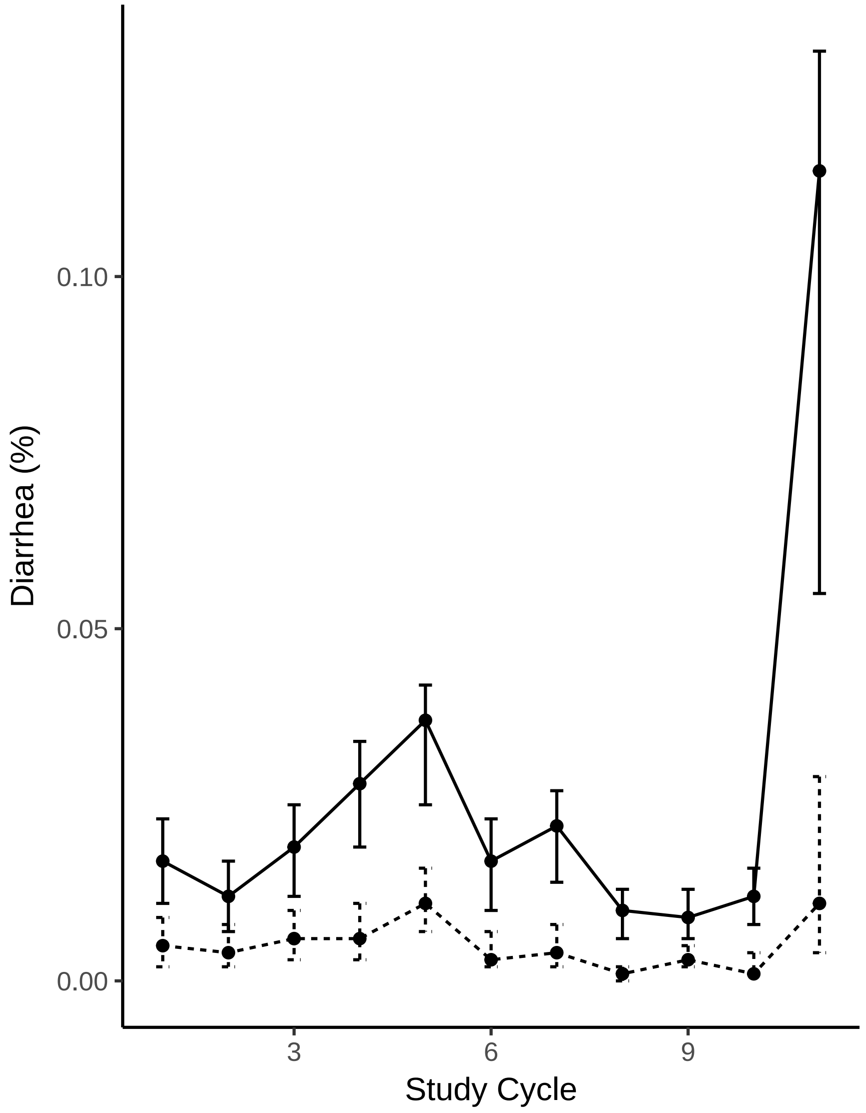


Figure S-2: Prevalence of diarrhea not attributable to rotavirus (solid) and attributable to rotavirus (dashed) over time

#### S3.2 Additional regression results

##### S3.2.1 Socioeconomic status

The socioeconomic indicators used here were not significant in the full population for either rotavirus infection or all-cause diarrhea, but had stronger associations for the youngest age group in the age-stratified

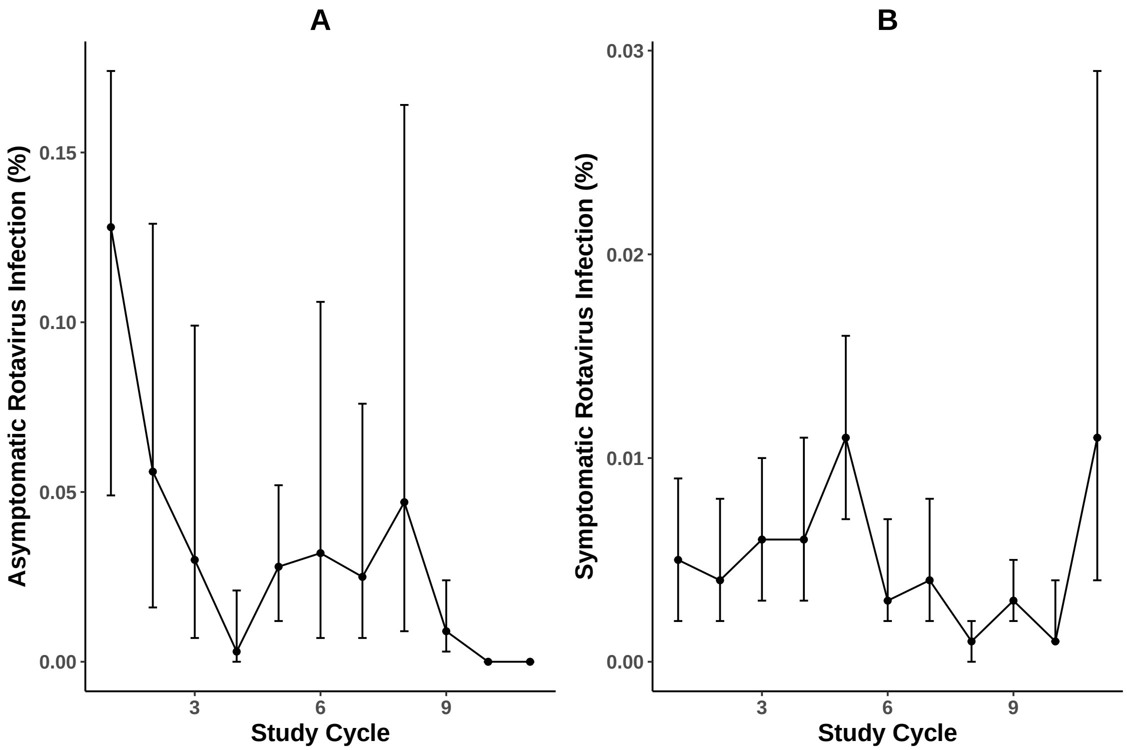


Figure S-3: Prevalence of A) Asymptomatic rotavirus infection and B) Symptomatic rotavirus infection over time. The rotavirus vaccine was introduced in study cycle 8

model. In general, higher household education and smaller household size would both be expected to indicate higher socioeconomic status.

For the youngest age group, increased household size was associated with increased diarrhea and rotavirus whereas it was protective among older age groups. For this age group, education was a significant predictor of rotavirus infection, with higher education being associated with decreased rotavirus infection. This association was not significant for all-cause diarrhea for any age group. Conceptually, this result is somewhat intuitive because the younger age groups with lower socioeconomic status would tend to get exposed earlier in life and might have increased protection as adults.

##### S3.2.2 All-cause diarrhea

The full regression table for the general population is shown in table S-6. After adjusting for relevant confounders, community vaccine coverage was no longer significantly associated with all-cause diarrhea in these six communities.

In this table, model 2 adjusts for rotavirus infection. In the main text (table 4), the vaccine efficacy estimate is not adjusted for whether or not rotavirus was a cause of that diarrhea in order to capture the full effect, given that we are not explicitly modeling synergy between rotavirus and co-infecting pathogens. The vaccine effectiveness estimates for all-cause diarrhea were extremely similar when adjusted for rotavirus infection, but were slightly attenuated and the effect estimates crossed the null.

| | Unadjusted | Model 1 | Model 2 |
|---|---|---|---|
| Rotavirus positive | 9.38 (5.95, 14.8) | 8.08 (4.15, 15.7) | 7.87 (4.04, 15.4) |
| Male | 1.12 (0.905, 1.40) | 1.09(0.784, 1.52) | 1.09 (0.782, 1.52) |
| Age | | | |
| $<1$ | 10.6 (6.95, 16.3) | 8.79(4.99, 15.0) | 8.78 (4.98, 15.5) |
| 1–5 | 12.8 (9.61, 16.9) | 13.2 (9.44, 18.4) | 13.2 (9.41, 18.4) |
| $\geq 5$ | Ref | Ref | Ref |
| Household size | 0.984 (0.949, 1.02) | 0.982 (0.931, 1.04) | 0.982 (0.931, 1.04) |
| Highest household education | 0.956 (0.926, 0.987) | 0.977 (0.934, 1.02) | 0.979 (0.936, 1.02) |
| Remoteness | | | |
| Close | Ref | Ref | Ref |
| Medium | 1.24 (0.841, 1.81) | 1.88 (1.15, 3.07) | 1.87 (1.14, 3.06) |
| Far | 0.775 (0.607, 0.990) | 1.18 (0.844, 1.66) | 1.21 (0.862, 1.70) |
| Community vaccine coverage | 0.602 (0.451, 0.803) | – | 0.848 (0.577, 1.25) |
| Vaccine introduced | 0.654 (0.512, 0.835) | – | – |

Table S-6: Vaccination coverage and all-cause diarrhea. All cells represent OR (95% CI). Odds ratios are calculated with respect to the reference group for categorical variables. Vaccine coverage is included as a proportion, taking a minimum value of 0 and a maximum value of 1. Model 1 is adjusted for rotavirus infection, gender, age, remoteness, household size, and highest household education. Model 2 is adjusted for all variables in Model 1 plus community vaccine coverage. Note that to calculate the percent reduction in diarrhea (Table 5 in the main text), we did not adjust for rotavirus infection and re-ran model 2 without adjusting for this variable.

| Age group | Unadjusted OR (95% CI) | Adjusted OR (95% CI) |
|---|---|---|
| <1 | 5.61 (1.24, 25.2) | 5.19 (1.16, 23.2) |
| 1-5 | 5.86 (1.87, 18.3) | 5.51 (1.74, 17.5) |
| ≥5 | 10.1 (5.37, 19.0) | 10.8 (5.85, 19.8) |
| Total population | 8.20 (4.94, 13.61) | 8.69 (4.31, 17.5) |

Table S-7: Effect of rotavirus infection on diarrhea illness by age. The adjusted odds ratios are adjusted for sex, household size, highest household education, and remoteness. Each odds ratio can be interpreted as the multiplicative increase in odds of diarrhea, given rotavirus infection.

###### S3.2.3 Rotavirus and diarrhea by age group

In order to determine if rotavirus was a causative diarrheal pathogen for all age groups, we also calculated the odds ratio for diarrhea given rotavirus infection relative to the odds of having diarrhea given no rotavirus infection. The results are shown in table S-7. Rotavirus is strongly associated with diarrheal symptoms in all age groups, including older children and adults (age $\geq 5$).

#### S4 Supplemental information for the cohort analysis

The regression results presented in the main text are also shown in Table S-8 to illustrate differences between the adjusted and unadjusted models. In this population, none of the socioeconomic indicators were significantly associated with all-cause diarrhea, unlike in the case control study. This difference is probably due to the fact that children with vaccine records were not comparable to children without vaccine records–children with vaccine records had lower household education than children without vaccine records.

While children who were older than 2 years of age had a lower rate of all-cause diarrhea, considering all episodes that occurred during the follow up period, they had a higher hazard of diarrhea. This difference reflects the fact that most diarrheal infections are incompletely immunizing. There as no evidence that the effect of vaccination was stronger among young children (based on the lack of significance of the interaction term).

#### S5 Assessment of seasonality

Chance confounding could have occurred if certain communities with particularly low/high levels of vaccine coverage were sampled for the case control study only in certain times of the year after the vaccine was introduced. To test for this possibility, we re-ran all models used to generate table 5 with and without adjusting for season as a binary variable (1 = January to May representing the rainy season and 0 = June to December representing the dry season). In these adjusted models, the associations between vaccine coverage and all-cause diarrhea and rotavirus infection changed by less than 5% for almost all age groups

Table S-8: Cox Regression Model for the time to first all-cause diarrhea episode among children. Models 2 and 3 are adjusted for household size (kids), highest household education, gender, community remoteness, and BCG vaccination.

| | Model 1<br>Unadjusted<br>HR (95% CI) | Model 2<br>Adjusted<br>HR (95% CI) | Model 3<br>Adjusted + Interactions<br>HR (95% CI) |
|---|---|---|---|
| Rotavirus Vaccine | | | |
| 0 doses | *Ref* | *Ref* | *Ref* |
| 1 dose | 1.02(0.552, 1.89) | 0.603 (0.290, 1.25) | 0.606 (0.269, 1.37) |
| 2 doses | 0.749(0.457, 1.23) | 0.474 (0.257, 0.874) | 0.451 (0.231, 0.882) |
| Age $\geq$2 years | 2.67(1.37, 5.22) | 2.35(1.19, 4.66) | 1.95 (0.393, 9.68) |
| Male | 0.843 (0.6, 1.18) | 0.914 (0.636, 1.31) | 0.913 (0.635, 1.31) |
| BCG Vaccination | 1.60 (0.706, 3.64) | 2.31 (0.906, 5.88) | 2.38 (0.914, 6.19) |
| Household Size | 1.01(0.917, 1.12) | 0.977 (0.881, 1.08) | 0.976 (0.878, 1.09) |
| Highest Household Education | 0.980 (0.930, 1.03) | 0.965 (0.913, 1.02) | 0.965 (0.914, 1.02) |
| Remoteness | | | |
| Close | *Ref* | *Ref* | *Ref* |
| Medium | 1.48 (0.818, 2.68) | 1.53 (0.830, 2.83) | 1.54 (0.835, 2.89) |
| Far | 2.03 (1.41, 2.91) | 2.28 (1.53, 3.40) | 2.28 (1.53, 3.40) |
| Age x Vaccine Interaction | | | |
| Age($\geq$2) x 1 dose | – | – | 1.06 (0.174, 6.45) |
| Age ($\geq$2) x 2 doses | – | – | 1.29 (0.259, 6.43) |

and outcomes. The two exceptions were both for children aged 1-5 years old. In this age group, the season-adjusted all-cause diarrhea effect estimate was 33.2% (95% CI: -20.7%-63.1%, a change of 9.7%) and the season-adjusted symptomatic rotavirus infection estimate was 44.4% (95% CI: -61.3%-80.8%, a, 10.3% absolute change). However, in both cases the main conclusions from the model remained the same.

### S6 Sensitivity analysis for coverage

The case-only and case-control analysis models which use community vaccine coverage as an exposure could be biased if community coverage was much lower or higher than expected. To address this concern, we re-ran all analyses for the case-control study both 1) assuming vaccine coverage was similar among children with and without vaccine records (the version in the main text) and 2) where vaccine coverage among children without records was set to a particular threshold of coverage (0%, 25%, 50%, 75%, and 100%). After assigning the new coverage level to children without vaccine records, we re-estimated the overall community level coverage using the following formula:

$$Coverage = p_{record}(Coverage_{record}) + (1 - p_{record})(Coverage_{norecord})$$

Where $p_{record}$ is the fraction of individuals with vaccine records, $Coverage_{record}$ is the coverage of two doses of Rotarix among children with vaccine records (based on data), and $Coverage_{norecord}$ is the coverage of two doses of Rotarix among children without vaccine records (varied systematically: equal to the community mean, 0%, 25%, 50%, 75% or 100%). We then re-ran the fully adjusted model with the new community vaccine coverage as the exposure. Results for rotavirus infection are shown in Figure S-6 and results for all-cause diarrhea are shown in Figure S-7 In general, results were highly similar for rotavirus infection for all coverage levels considered and the point estimates were in fact stronger when vaccine coverage among children without records was lower than the community average. When the equal coverage assumption was used, the effect was not statistically significant for children $<$1 year of age but the confidence intervals were narrower and became statistically significant when other assumptions were used. The all-cause diarrhea estimates were more variable with different coverage assumptions, but nearly all associations remained null. The one exception was the estimate for children $<$1 year, which had a significant protective effect when coverage for children without vaccine records was 100%.

### S7 Internal consistency of results between analysis parts

Prior to doing any analysis, we decided which communities would be included in each part of the analysis in an effort to minimize bias in our vaccine effectiveness estimates. Because we used different communities for different parts of this analysis, we compared the results between them to ensure that our analysis was generalizable to all communities across the region and that the associations we found were not purely the result of our sampling process. In general, our results were highly internally consistent throughout the region.

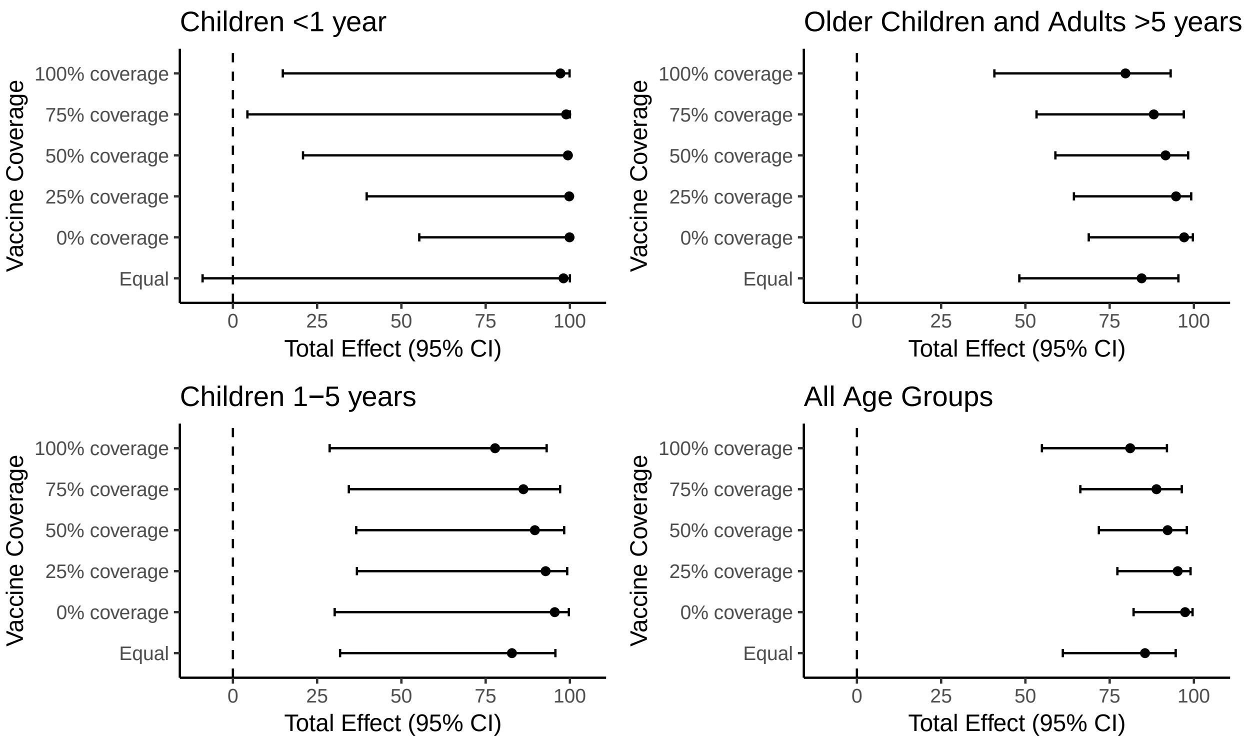


Figure S-4: Total effect of vaccination (100% vs. 0% coverage) on rotavirus infection (part 1 analysis) by level of vaccine coverage among children without records (0%, 25%, 50%, 75%, 100% or equal to the community average).

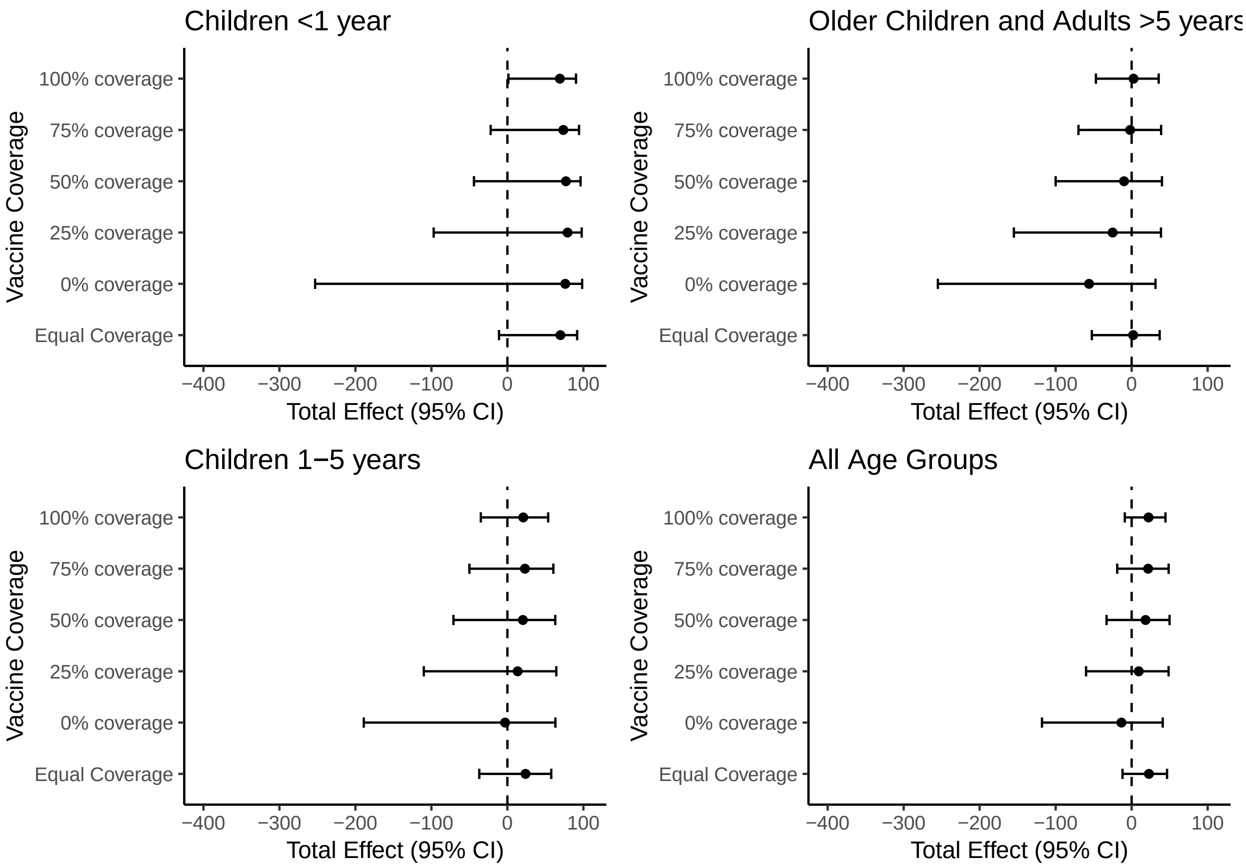


Figure S-5: Total effect of vaccination (100% vs. 0% coverage) on all-cause diarrhea (part 1 analysis) by level of vaccine coverage among children without records (0%, 25%, 50%, 75%, 100% or equal to the community average).

To compare parts 1 and 3, we subset the data from the overall case-control study and compared the results only among children less than five years of age. Because the all-cause diarrhea estimates from the case control study were highly similar both before and after adjusting for vaccine coverage (i.e., the covariates and vaccine coverage were independent predictors of all-cause diarrhea, see Table S-6) we included all study years in this comparison, including time both before and after the vaccine was introduced. By design, the odds ratio approximates the rate ratio (from the Poisson sensitivity analysis) in the case control study because controls were time matched to cases and because the outcome was rare in all exposure groups. The results are shown in table S-9.

The vaccine efficacy estimate was higher in the cohort analysis. This difference is consistent with other published data showing that studies investigating the rate of all-cause diarrhea rather than the hazard may tend to underestimate vaccine effectiveness [3]. Consistent with this body of work, we found that older children had a higher hazard of all-cause diarrhea but a lower rate (estimated using Poisson regression, IRR=0.503, 95% CI: 0.361, 0.700). Using Poisson regression also produced an effect estimate of 39.8% for children under 2, which is within 7% of the estimate calculated using case control data and is similar to results presented in vaccine trials, which estimated 39-42% [4, 5]. Older children who had an all-cause diarrhea episode tended to have fewer episodes than younger children.

Another difference is the association between remoteness and all-cause diarrhea. In the case control analysis, far communities had similar rates of disease to close communities whereas in the cohort analysis, far communities were at higher risk of disease. This difference most likely arose because children with vaccine records tended to come disproportionately from close/medium villages. Therefore, it is likely that we are not estimating the rate ratio well for far communities in the cohort analysis. The association with education was also slightly different in the cohort analysis, which is also consistent with the marginally significant association we found comparing children with vaccine records to children without vaccine records.

| | Case-Control Analysis (Part 1) OR (95% CI) | Cohort Analysis-Cox (Part 3) HR (95% CI) | Cohort Analysis-Poisson (sensitivity) IRR (95% CI) |
|---|---|---|---|
| Male | 1.05 (0.686, 1.61) | 0.878(0.612, 1.26) | 0.983 (0.764, 1.27) |
| Age | | | |
| <2 | Ref | Ref | Ref |
| 2–5 | 0.426 (0.277, 0.657) | 2.23 (1.13, 4.42) | 0.502 (0.359, 0.7) |
| Household size | 0.973 (0.900, 1.05) | 0.995 (0.942, 1.05) | 0.992 (0.955, 1.03) |
| Highest household education | 0.941 (0.881, 1.01) | 0.976 (0.924, 1.03) | 1.01 (0.966, 1.03) |
| Remoteness | | | |
| Close | Ref | Ref | Ref |
| Medium | 2.07 (0.925, 4.62) | 1.42 (0.771, 2.60) | 1.29 (0.852, 1.96) |
| Far | 1.15 (0.693, 1.91) | 2.18 (1.48, 3.22) | 1.42 (1.08, 1.86) |
| Effect of 2 doses* | 46.3% (-22.8%, 76.5%) | 57.1% (16.6%-77.9%) | 39.8% (13.7%, 58.0%) |

Table S-9: Comparison between parts 1 and 2. All parameters except the effect of two doses were calculated using all children in the case-control dataset under 5 years of age for part 2 and all children in the cohort of part 1 and were estimated in a single model. For the effect of two doses, we subset both data files at 2 years of age and presented calculated vaccine effectiveness using the overall coverage parameter for the case control analysis and the two doses for the cohort analysis.
